# Ligand-responsive groove remodelling in human and macaque CD1d reveals a conserved MHC-like gating mechanism

**DOI:** 10.64898/2025.12.23.695970

**Authors:** Daniel Burns, Alex Look, Steven Turner, Martin Malý, Oliver Melling, Kinga Niedobecka, Rita Szoke-Kovacs, H. Nurdan Aksoy Kilinc, Richard Suckling, Andrew Chancellor, Mariolina Salio, Andrew White, Sally Sharpe, Ali Roghanian, Bruno Linclau, Paul Elkington, Jonathan W Essex, Ivo Tews, Salah Mansour

## Abstract

CD1d presents lipid antigens to invariant natural killer T (iNKT) cells. We determined a high-resolution crystal structure of human CD1d bound to α-galactosylceramide (α-GalCer) at 1.76 Å, enabling detailed investigation of ligand-sensitive conformational flexibility at Phe84, a conserved aromatic residue that caps the F′ groove. Electron density at Phe84 revealed multiple side-chain conformations, suggestive of ligand-induced plasticity. Molecular dynamics simulations indicated that the canonical rotamer is energetically favoured in the absence of a stabilising groove-occupying ligand. To assess conservation of this putative gating mechanism, we solved the first CD1d structure from a non-human primate, rhesus macaque CD1d-α-GalCer, at 1.83 Å resolution. In contrast to the human complex, Phe84 in macaque CD1d adopted a fixed conformation. As this aromatic residue is conserved across CD1 isoforms and CD1d-expressing species, and mirrors gating residues in MHC class I that regulate peptide accommodation, our findings support a shared evolutionary strategy for managing antigen diversity. These data provide critical insight into the mechanisms of antigen presentation by CD1 molecules.

**Significance Statement:** This study reveals that Phe84, a conserved aromatic residue in CD1d, may act as a ligand-responsive gate modulating F′ groove accessibility. This conditional plasticity could enable binding of structurally diverse lipid antigens and appears conserved across CD1 isoforms. The mechanism parallels class I MHC, where gating residues regulate peptide presentation, suggesting an evolutionarily shared strategy for accommodating antigen diversity.

## Introduction

CD1 proteins represent a lineage of antigen-presenting molecules distinct from HLA and MR1 (1, 2). Encoded on chromosome 1, CD1 molecules are classified into three groups based on sequence homology. CD1 isoforms differ in cellular expression, intracellular trafficking, and lipid-binding capabilities, with the latter being dictated predominantly by the structural architecture of the antigen-binding groove (1, 3). Group 1 consists of CD1a, CD1b, and CD1c, while Group 2 comprises CD1d (1, 2). Group 3 includes CD1e, which does not localise to the cell surface but instead functions as an intracellular chaperone involved in lipid antigen processing (4, 5). CD1 molecules feature two major antigen-binding pockets, known as the A’ and F’ grooves. The A’ groove is generally more conserved across CD1 isoforms, whereas the F’ groove displays greater variability (6).

Structural studies of all four antigen-presenting human CD1 isoforms have mapped multiple lipid-binding interactions, providing mechanistic insight into their similarities and differences (7–10). Notably, CD1a has a wide and shallow F’ groove, optimised for accommodating lipopeptides such as didehydroxymycobactin (8). CD1b, in contrast, possesses a unique portal connecting the A’ and F’ grooves, thereby extending the overall groove length and enabling the binding of exceptionally long acyl chains, such as those found in mycobacterial mycolates (7, 11). Structural characterisation of CD1c further revealed that the F′ groove can adopt distinct open and closed conformations, underpinned by a ligand-sensitive histidine side-chain double-conformation, thereby permitting accommodation of diverse lipid species (9, 12, 13).

CD1d is functionally distinct in that it is recognised by invariant natural killer T (iNKT) cells, a specialised T cell lineage with potent anti-tumour activity (14). Upon activation by lipid antigens presented by CD1d, iNKT cells rapidly secrete cytokines, such as interferon-gamma (IFN-γ), that enhance anti-tumour immunity by activating natural killer cells, dendritic cells, and conventional T cells (14–17). This ability to bridge innate and adaptive immunity has made iNKT cells attractive candidates for cancer immunotherapy. Although murine models have provided fundamental insights into CD1d-restricted immunity, their translational relevance is limited by the absence of group 1 CD1 molecules (18). In contrast, non-human primates such as rhesus macaques express the full CD1 repertoire and display closer immunogenetic homology to humans (19). Macaques therefore offer a more physiologically relevant system for studying CD1d antigen presentation and iNKT cell activation. However, no high-resolution structural data exist for macaque CD1d, limiting cross-species comparisons.

Despite the well-characterised structure of human CD1d (10), key questions remain about how it accommodates large or atypical lipid antigens (20, 21). CD1d lacks the accessory portals seen in other CD1 isoforms, such as CD1b, yet it binds ligands with alkyl chains that exceed the calculated groove capacity (20). This suggests CD1d may instead rely on internal structural plasticity to accommodate larger lipids. CD1c’s F′ groove exhibits ligand-sensitive flexibility, adopting distinct conformations depending on antigen occupancy (9, 12, 13), but it remains unclear whether similar flexibility exists in CD1d. We hypothesised that CD1d may undergo conformational rearrangements within the groove, with Phe84, a conserved aromatic residue that caps the F′ groove, as a key gating element. This residue is analogous to Tyr84 in class I MHC, which regulates peptide accommodation through rotameric repositioning (22–24), and may play a similar role in modulating groove accessibility in CD1d.

Here, we present high-resolution crystal structures of human and macaque CD1d bound to α-GalCer. These structures enabled us to investigate the conformational behaviour of Phe84. While the crystallographic data revealed unexplained electron density near this residue in human CD1d, molecular dynamics simulations suggest that alternative rotamers are only transiently stable and likely require stabilisation by groove-occupying ligands. In contrast, the macaque CD1d structure adopts a single, canonical conformation at this position. These findings suggest that conformational plasticity at the F′ groove may be context-dependent, potentially contributing to a conserved gating mechanism that facilitates the accommodation of structurally diverse lipid antigens.

## Results

### The CD1d F’ groove in open and closed states

To investigate CD1d groove architecture, we determined a high-resolution crystal structure of human CD1d in complex with the potent iNKT agonist α-GalCer. Using established methods (7, 25, 26), we generated soluble CD1d-β2m complexes and solved the structure at 1.76 Å resolution (Figure 1, Table 1). This improves upon the previously published CD1d-α-GalCer structure (PDB: 1ZT4, 3.0 Å (10)), allowing more detailed modelling of side chains and lipid interactions.

**Fig. 1.**
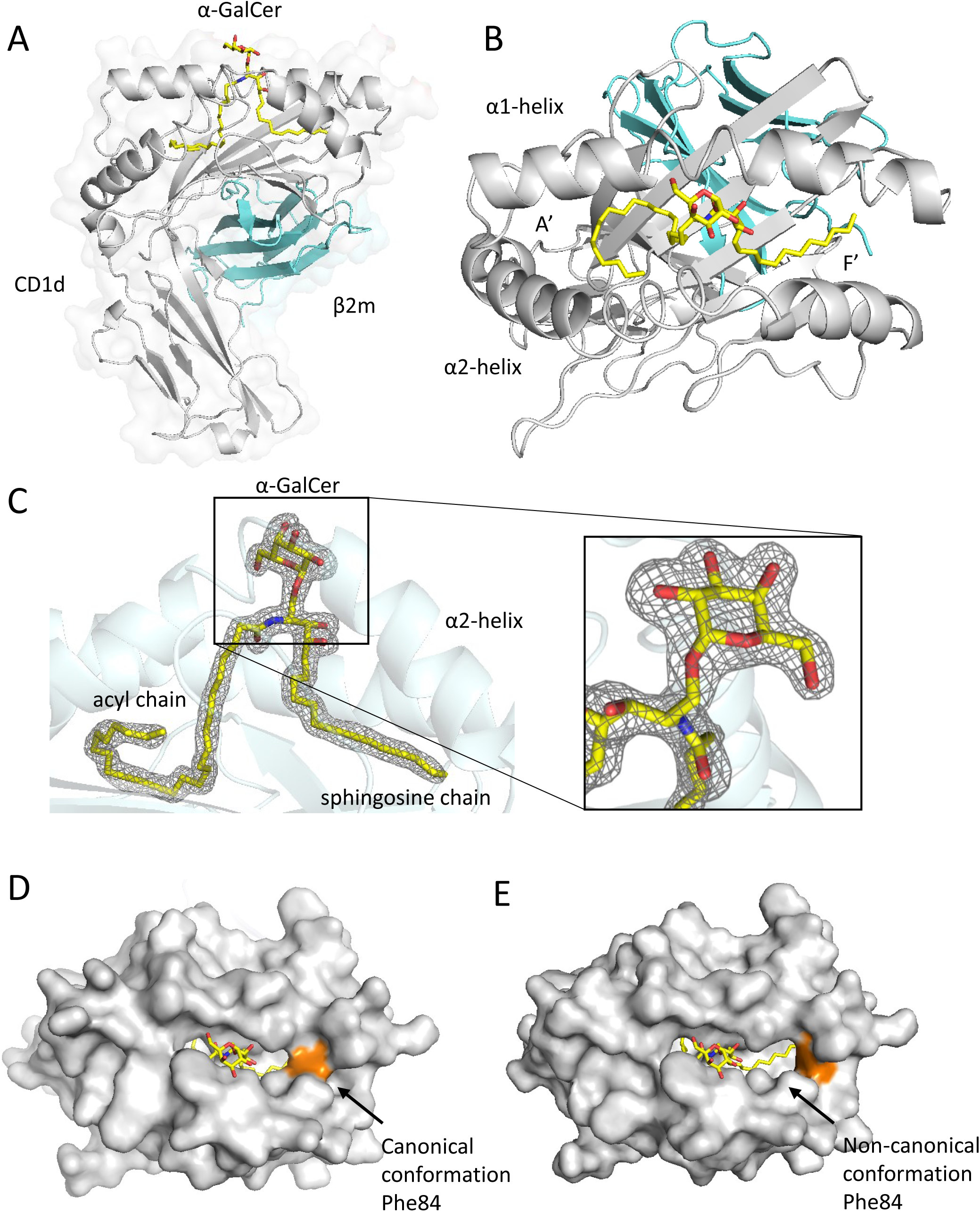
Human CD1d adopts alternative F′ groove conformations via Phe84. **A)** Overall structure of human CD1d-β2m bound to α-GalCer, showing the galactose head group exposed at the surface and lipid tails buried within the A′ and F′ channels. **B)** Top-down view highlighting α-GalCer’s acyl and sphingosine chains occupying the A′ and F′ grooves, respectively. **C)** Omit map contoured at 3σ confirms well-resolved α-GalCer density. **(D-E)** Surface representations of CD1d in the **(D)** canonical and **(E)** non-canonical conformations of Phe84. Dual rotamers reshape the F′ roof, suggesting a flexible gating mechanism. Phe84 is shown in orange; CD1d in grey; β2m in cyan; α-GalCer in yellow (heteroatoms: N, blue; O, red).

**Table 1:** Data-collection and merging statistics and structure-refinement parameters for the crystal structures of Human and Macaque CD1d. For data processing statistics, values in parentheses are for the highest resolution shell. *R*1 is the *R*-value calculated from the square root of intensities where *I*/*σ*(*I*) > 2. Average CC is the correlation coefficient between calculated and observed intensities averaged over resolution shells. R.m.s.d. and R.m.s.Z. stand for root mean square deviation and Z score, respectively.

|  | Human CD1d | Macaque CD1d |
| --- | --- | --- |
| PDB entry | 9RSE | 9RSF |
| *Data collection* |  |  |
| X-ray source | ID30B, ESRF | I03, Diamond Light Source |
| Wavelength (Å) | 0.9253 | 0.9763 |
| Detector | DECTRIS PILATUS 6M | DECTRIS EIGER2 XE 16M |
| Temperature (K) | 100 | 100 |
| Crystal-to-detector distance (mm) | 371.2 | 326.6 |
| No. of oscillation images | 1800 | 3600 |
| Exposure time per image (s) | 0.02 | 0.007 |
| Oscillation width (°) | 0.05 | 0.1 |
| Transmission (%) | 1 | 10 |
| *Data processing* |  |  |
| Space group | $P2_12_12$ | $P3_221$ |
| a, b, c (Å) | 117.62, 156.37, 51.91 | 87.06, 87.06, 232.8 |
| Resolution (Å) | 49.27–1.76 (1.79–1.76) | 54.08–1.83 (1.86–1.83) |
| No. of reflections | 308,847 (10,565) | 1,525,765 (25,554) |
| No. of unique reflections | 92,489 (3,363) | 89,434 (3,817) |
| Multiplicity | 3.3 (3.1) | 17.1 (6.7) |
| Completeness (%) | 96.6 (71.1) | 98.2 (84.73) |
| $R_{\text{meas}}$ | 0.094 (0.937) | 0.056 (0.486) |
| $R_{\text{p.i.m.}}$ | 0.049 (0.487) | 0.013 (0.179) |
| Average $I/\sigma(I)$ | 14.8 (1.1) | 32.7 (2.1) |
| CC<sub>1/2</sub> | 0.968 (0.532) | 1.000 (0.914) |
| CC* | 0.992 (0.833) | 1.000 (0.977) |
| Wilson B (Å<sup>2</sup>) | 24.95 | 24.77 |
| Solvent content (%) | 56.1 | 58.6 |
| *Structure refinement* |  |  |
| $R_{\text{work}}$ | 0.2132 | 0.1710 |
| $R_{\text{free}}$ (2.2% reflections) | 0.2542 | 0.2171 |
| $R1_{\text{work}}$ | 0.1789 | 0.1611 |
| $R1_{\text{free}}$ | 0.22 | 0.2039 |
| Average $CC_{\text{work}}$ | 0.8668 | 0.9398 |
| Average $CC_{\text{free}}$ | 0.7682 | 0.8948 |
| Median ADP (Å) | 32.4 | 28.6 |
| No. of protein heterodimers in asymmetric unit | 2 | 2 |
| No. of non-hydrogen atoms | 7,225 | 7,438 |
| No. of water molecules | 710 | 1,077 |
| Ligands | 2 × α-Galactosylceramide, 2 × pentaethylene glycol | 2 × α-Galactosylceramide, 7 × ethylene glycol, 3 × Cl<sup>-</sup> |
| R.m.s.d. (R.m.s.Z.) from ideal bond lengths (Å) | 0.013 (0.99) | 0.014 (1.07) |
| R.m.s.d. (R.m.s.Z.) from ideal angles (°) | 2.08 (1.08) | 2.03 (1.09) |
| Ramachandran favoured (%) | 98.0 | 98.1 |
| Ramachandran outliers (%) | 0 | 0.3 |
| Rama-Z score | -0.14 ± 0.27 | -0.05 ± 0.28 |
| Clashscore | 5.8 | 3.5 |

CD1d adopted the expected MHC-like fold, with α-GalCer’s acyl and sphingosine chains fully occupying the A′ and F′ grooves, respectively. The galactose head group was positioned at the surface for potential TCR engagement (Figure 1A-C). The lipid was well-resolved, and functional assays confirmed robust iNKT binding and cytokine responses (Supplementary Figure 1A-E).

We observed additional electron density above the F′ groove, between α-GalCer and the side chain of Phe84, a conserved residue that caps the groove. This density could not be fully accounted for by the ligand or protein alone and may reflect a disordered buffer molecule or minor occupancy by a stabilising moiety (Supplementary Figure 2). To explore potential conformational flexibility at this site, we modelled Phe84 in both canonical (inward) and non-canonical (outward) rotamers. Omit maps showed residual difference density for each individual rotamer, which was largely resolved when modelled as a dual conformation (Supplementary Figure 2A-D). These models suggest that Phe84 may sample alternate positions under specific conditions. Although omit maps suggested alternate positions, steric constraints indicate that the canonical (inward) rotamer is energetically preferred, and this was used for deposition. Nonetheless, the additional density suggests Phe84 may retain some conformational flexibility under specific conditions.

Surface representations of CD1d incorporating each Phe84 orientation highlight the impact of this side chain on F′ groove shape (Figure 1D-E). While the structural data alone do not demonstrate dynamic switching, they suggest a latent plasticity at this site. This echoes prior observations in CD1c, where conformational variability at the equivalent His84 residue modulates groove architecture (9). Further, the analogous position of Tyr84 in class I MHC (23, 27, 28) links to rotamer selection to regulate peptide accommodation (Supplementary Figure 3). From these previous observations, we hypothesised that gating at position 84 may be a conserved mechanism across CD1 isoforms and MHC-I.

### Aromatic gating residue conserved across the human CD1 isoforms

We analysed the sequences and published structural data of the four human CD1 isoforms to investigate the role of residues covering the F’ groove. Sequence analysis revealed conserved aromatic residues aligning with Phe84 in CD1d (Figure 2A): Tyr84 in CD1a (8), Phe84 in CD1b (7), and His84 in CD1c (9). We further detect conservation of alanine C-terminal to the F’ groove aromatic amino acids in all four CD1 isoforms. This conserved motif may suggest a prominent function, where the presence of a small residue following the aromatic amino acid may provide some structural flexibility (Figure 2A). The hydrophobic side chain sequesters the CD1 groove from solvent (Figure 2B; highlighted in orange). We next sought to determine whether the aromatic amino acid exhibits conformational plasticity across CD1 isoforms and could provide a gating mechanism, by analysing available structural data.

**Fig. 2.**
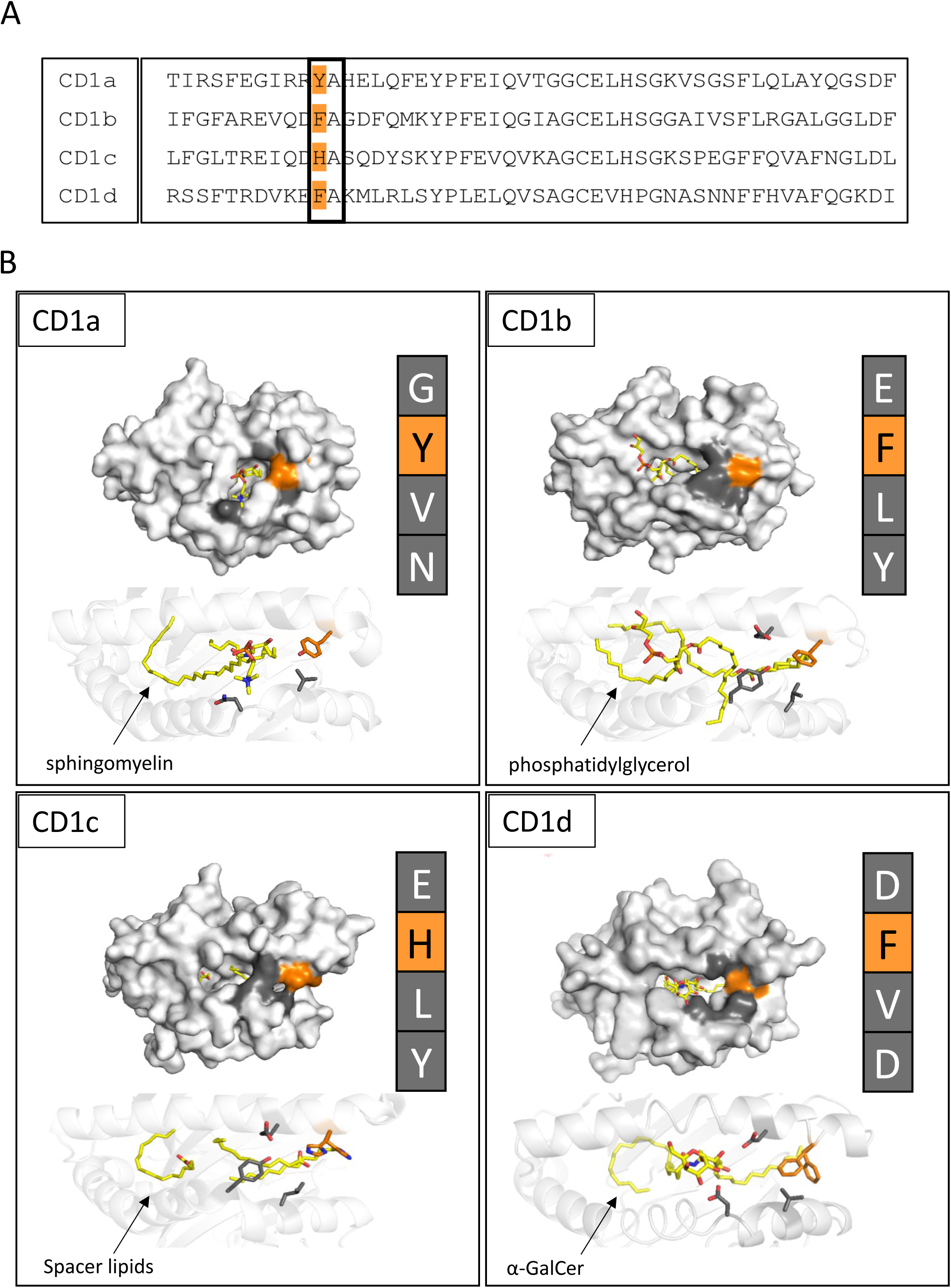
Aromatic gating residues are conserved across CD1 isoforms. **A)** Sequence alignment of human CD1a-d reveals conserved aromatic residues (Tyr or Phe) at position 84. **B)** Structural comparison of CD1 isoforms (CD1a, CD1b, CD1c, CD1d) showing location and orientation of residue 84 above the F′ groove. Surface and cartoon views highlight the potential for structural flexibility across isoforms. PDB codes 4X6F, 5WL1, 5C9J, and 1ZT4 used for CD1a, CD1b, CD1c, and CD1d respectively.

### Amino acid 84 displays multiple conformations on human CD1 isoforms

Several human CD1 crystal structures suggest flexibility at position 84, which sits above the F′ groove. In CD1b, Phe84 adopts distinct conformations depending on the lipid ligand, as seen in CD1b-GM2 and CD1b-PC complexes (7, 29) (Figure 3A). These alternate side chain orientations resemble the dual Phe84 conformations we modelled in CD1d (Figure 3A). Similarly, the equivalent residue in CD1c-His84 in CD1c-MPM (12) and His87 in CD1c-SL (9), also exhibits conformational variability, resulting in open or closed F′ groove arrangements (Figure 3B). Superimposing these structures with human CD1d highlights shared positional flexibility, supporting the idea that aromatic gating at this site may be a conserved feature within the CD1 family (Figure 3B). Having identified alternative F′ groove conformations across human CD1 isoforms, we next explored whether this plasticity is conserved across species.

**Fig. 3.**
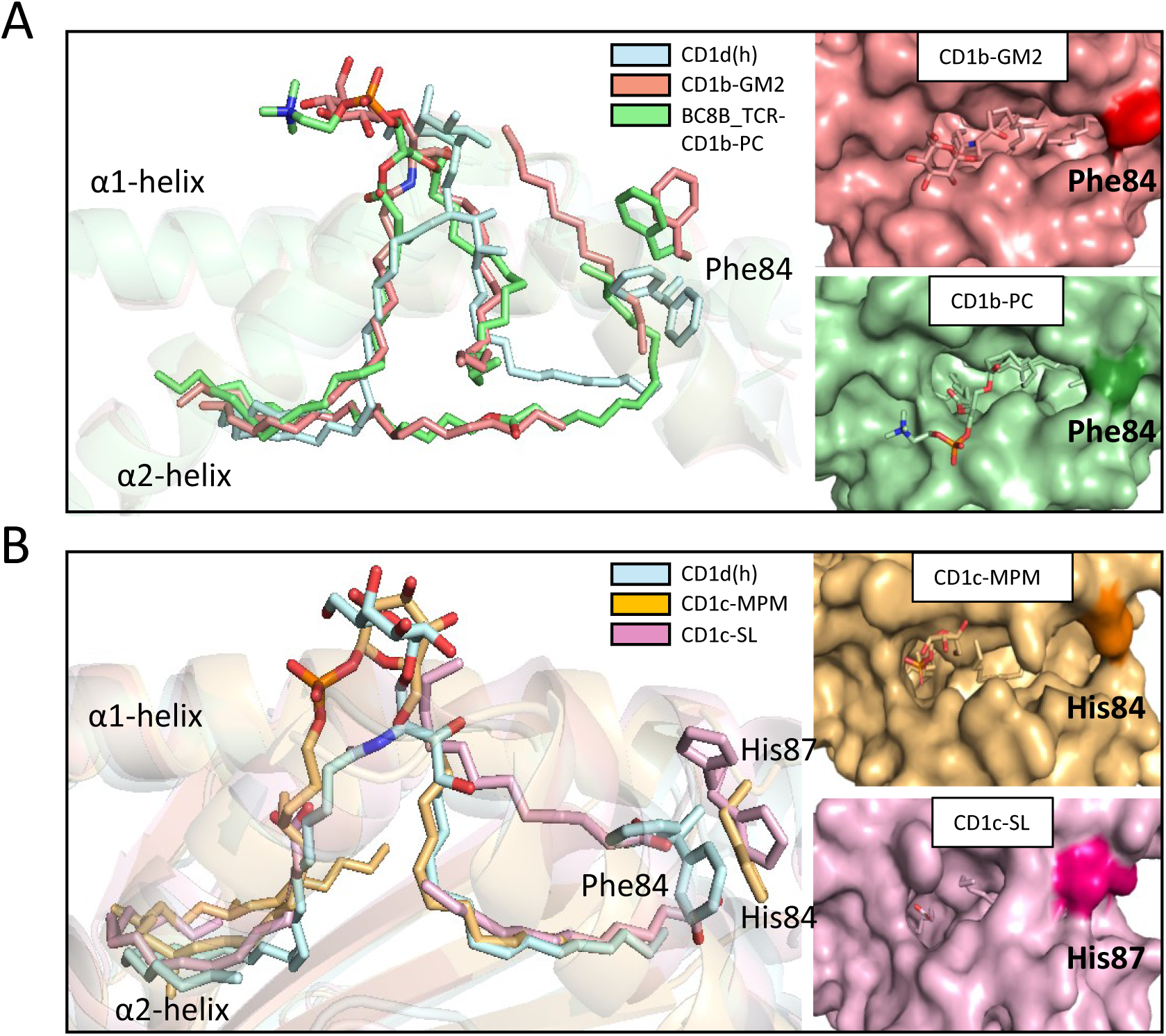
Conformational flexibility at residue 84 is shared among CD1 isoforms. **A)** Overlay of human CD1d (dual Phe84 conformations) with CD1b-GM2 and CD1b-PC shows distinct rotamer states associated with ligand occupancy. **B)** Similar comparisons with CD1c-MPM and CD1c-SL reveal alternative conformations of His84/His87, consistent with an open or closed F′ groove. These data support a conserved gating mechanism mediated by aromatic residues in CD1b, CD1c, and CD1d. Lipids are shown in stick representation with nitrogen, oxygen, and carbon shown in blue, red, and yellow respectively. Human CD1d, light blue; CD1b-GM2, dark pink; CD1b-PC, green; CD1c-MPM, yellow; CD1c-SL, pink. PDB codes 6CUG for CD1b-PC, 1GZP for CD1b-GM2, 5C9J for CD1c-SL, and 3OV6 for CD1c-MPM.

### Phe84 is conserved on CD1d across species

Sequence conservation across multiple species, including macaques, supports a conserved functional role for Phe84 in CD1d-expressing lineages (Figure 4A). However, rodent-like species do not retain phenylalanine at this position. Structural analysis of available CD1d molecules from human (10), bovine (30), and mouse (31) reveals that human and bovine CD1d are highly similar, with Phe84 adopting a canonical conformation that shields the sphingosine chain of α-GalCer from solvent (Figure 4B).

**Fig. 4.**
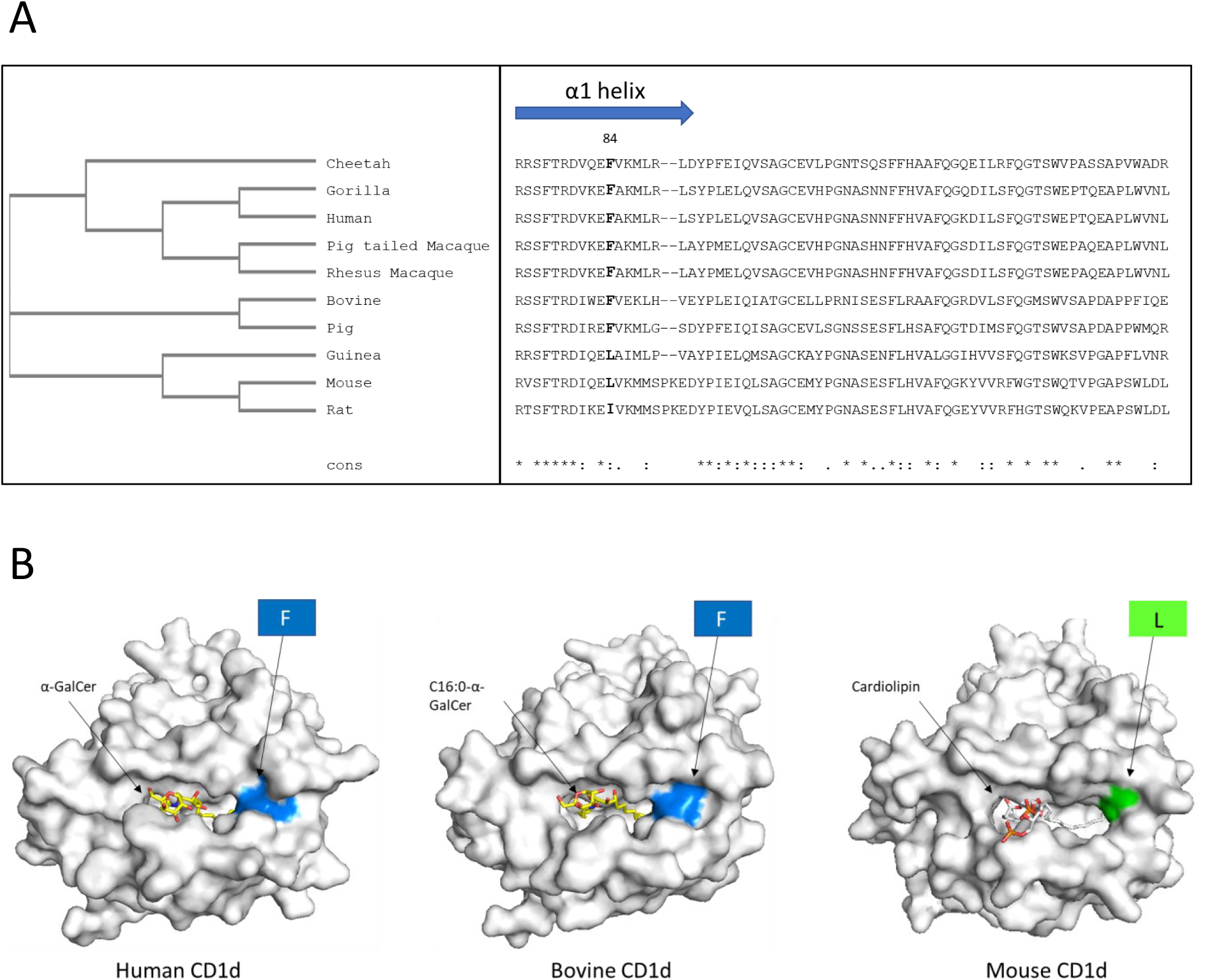
Cross species conservation of Phe84. **A)** Phylogenetic tree, sequence alignment and percentage identity of CD1d from 10 CD1d expressing species. Phe84 is conserved across the majority of species except rats, mice, and Guinea pigs. **B)** Comparison of current published structural data of CD1d from human, cow, and mouse. Phe84 highlighted in blue. The equivalent residue Leucine 84 in mouse highlighted in green. CD1d, grey; lipid molecules, yellow and coloured by heteroatom. PDB codes 1ZT4, 4F7E, 3MA7 from left to right respectively.

In contrast, mouse CD1d encodes isoleucine at this position (31), which lacks an aromatic ring and results in an ‘open’ conformation. This structural difference expands the F′ groove and exposes the binding cavity more extensively than in human and bovine CD1d (Figure 4B). These findings reveal fundamental molecular differences between rodent CD1d and that of higher species. While this suggests evolutionary divergence in rodents, the conservation of Phe84 in primates led us to investigate the structural properties of CD1d in rhesus macaques, a physiologically relevant model species.

### Crystal structure of macaque CD1d-α-GalCer

Macaques are valuable models for studying human disease. Functionally, macaque CD1d tetramers stained human iNKT cells strongly (Supplementary Figure 1A-B), induced iNKT activation and IL-2 production in a dose-dependent manner (Supplementary Figure 1C-D), and demonstrated direct TCR binding by SPR, with no cross-reactivity to CD1c (Supplementary Figure 1E).

To investigate its structure, we crystallised rhesus macaque CD1d in complex with α-GalCer, generating soluble CD1d/β2m proteins. The structure, refined at 1.83 Å resolution (Figure 5, Table 1), closely resembled human CD1d, with an RMSD of 0.77Å over 372 Cα atoms. The acyl and sphingosine chains were deeply buried within the A′ and F′ grooves, and the galactose head group was clearly resolved at the solvent-exposed surface (Figure 5A-B).

**Fig. 5.**
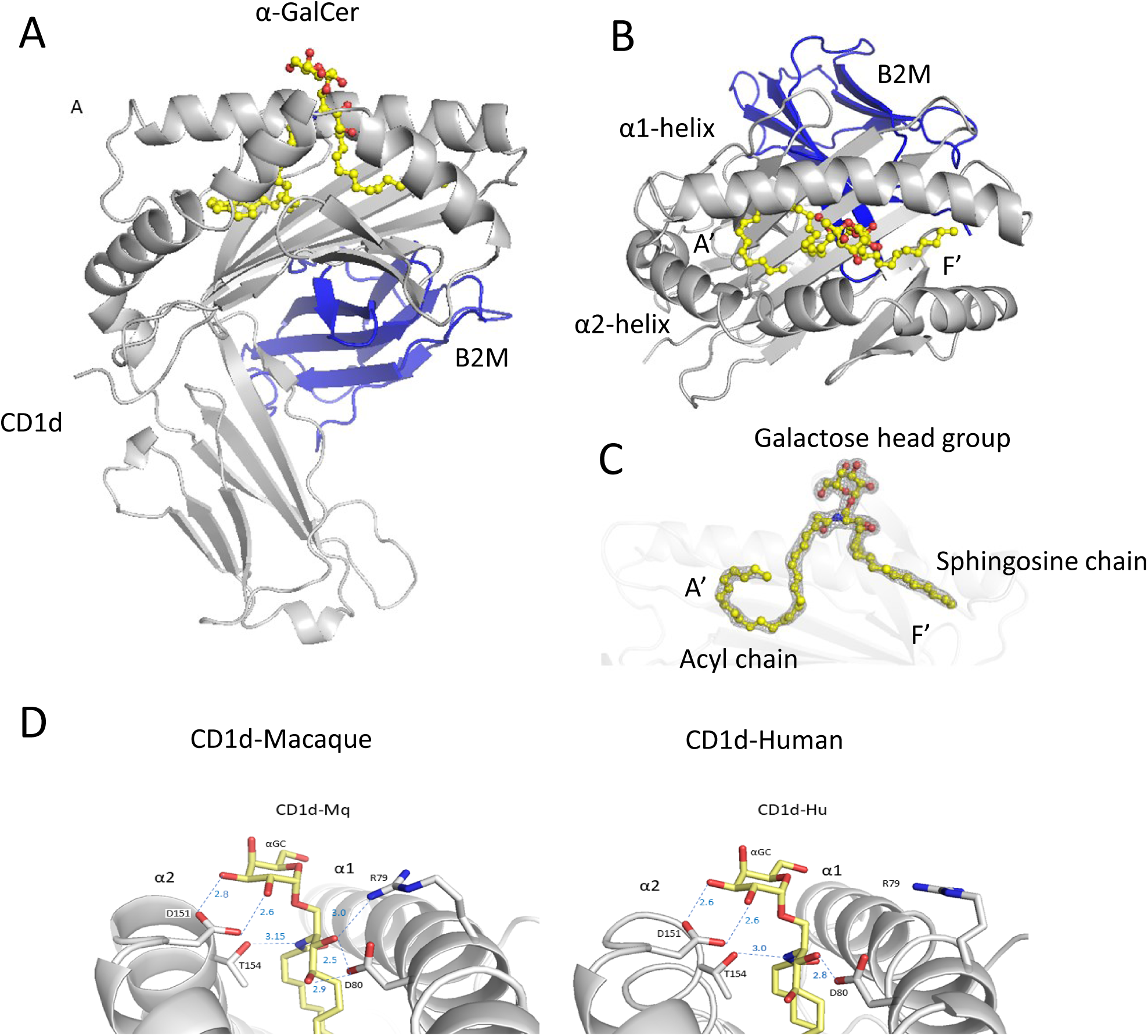
Overview of rhesus macaque CD1d-α-GalCer structure. (A-B) Crystal structure of macaque CD1d-β2m bound to α-GalCer shows conserved overall fold and lipid positioning within A′ and F′ grooves. **C)** Electron density confirms lipid occupancy. **D)** Comparison of the protein-lipid hydrogen bonding network between human and macaque CD1d. Hydrogen bonds to key amino acids shown in blue with distances labelled. Reorientation of Arg79 prevents it from the hydrogen bonding network in human CD1d. CD1d heavy chain, grey; β2m, dark blue; α-GalCer, yellow and coloured by heteroatom with carbon in yellow, oxygen in red, and nitrogen in blue.

**Fig. 6.**
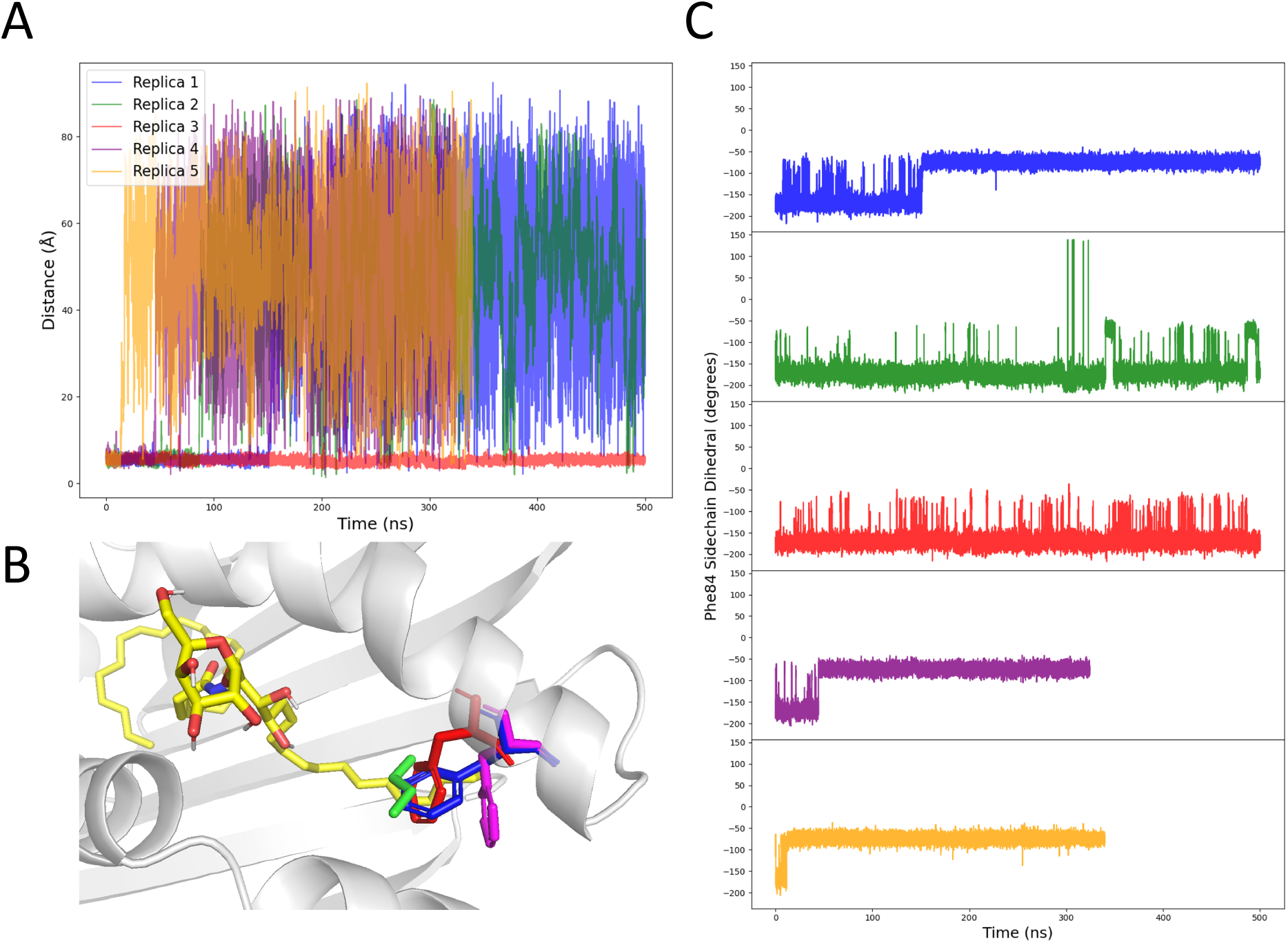
Ligand-dependent stabilisation of Phe84 conformations in human CD1d. **A)** Distance between α-GalCer and butane (centre of mass) over simulation time, showing butane dissociation for all five replicas. **B)** Representative structures from MD simulations showing Phe84 in non-canonical conformation with butane bound (red) and canonical state with butane unbound (blue). Representative conformations of α-GalCer (Yellow) and butane (Green) are overlaid for reference **C)** Phe84 dihedral angle (N–Cα–Cβ–Cγ) across simulations. Presence of butane stabilises the non-canonical rotamer, supporting ligand-induced gating behaviour.

Interestingly, we observed subtle differences in the lipid-binding groove. Relative to human CD1d, Arg79 on the α1 helix adopts a rotated conformation in macaque CD1d, disrupting the hydrogen bond to the ceramide C2 oxygen of α-GalCer chain (Figure 5C). In contrast to human CD1d (10), macaque CD1d exhibited no additional electron density above the F′ groove near Phe84. The lipid-binding groove appeared well-ordered, though this does not exclude the possibility that conformational changes may occur in response to specific ligands.

In the α-GalCer-bound macaque CD1d structure, Phe84 remained in the canonical (closed) conformation, restricting the width of the F′ groove. While we did not observe conformational switching in the macaque structure, we used molecular dynamics (MD) simulations of human CD1d to test whether Phe84 exhibits ligand-sensitive flexibility, particularly under conditions where electron density suggested alternative side-chain positions.

### Molecular dynamics simulations of human CD1d

Crystallographic density maps suggested the possibility of alternative rotamer conformations for the Phe84 side chain, potentially enabling ligand gating in human CD1d. To investigate this, we performed two REST2 MD simulations, each initiated from a different starting conformation of Phe84. These conformations were distinguished by the dihedral angle defined by the N–Cα–Cβ–Cγ atoms: the canonical form at −84.4° and the non-canonical form at −161.1°.

Across both REST2 MD simulations, the canonical form was energetically favoured. When the simulation was initiated with the non-canonical conformation, Phe84 rapidly rotated into the canonical position within 20 ps of NVT equilibration (Supplementary Figure 4A). The subsequent REST2 trajectory did not sample the non-canonical conformation (Supplementary Figure 4B) during 200 ns of simulation, indicating that the non-canonical rotamer is unstable in the context of CD1d bound to α-GalCer alone.

The REST2 simulation that began with Phe84 in the canonical form sampled both the canonical and non-canonical conformations with 0.67 % of frames occupying the non-canonical conformation. From the relative populations of the two states, we can estimate the free energy difference to be approximately 3 kcal mol^-1^.

However, the electron density maps from our crystallographic data showed additional unresolved density between α-GalCer and Phe84, raising the possibility that the non-canonical state might be stabilised by an unidentified co-factor, such as a crystallisation reagent. To explore this, we placed a hydrophobic four-carbon spacer (butane) into this region and simulated the system with the Phe84 starting in the non-canonical conformation. We chose butane as a minimal hydrophobic spacer ligand, reflecting the short-chain aliphatic lipids frequently observed as ‘spacer’ ligands in CD1 grooves (20). We then performed five independent conventional MD simulations, each of 500 ns in length. In all five repeats, the non-canonical conformation was clearly the most populated state, suggesting it was stabilised by the presence of the butane. In four of the five repeats the butane unbound resulting in the Phe84 adopting the canonical conformation in three of these four simulations. In the fourth simulation during which the butane unbound the sidechain angle of Phe84 did not return to the canonical conformation. A displaced conformation was observed, in which the backbone had shifted such that the Phe84 sidechain was now occupying the space that had previously been occupied by the butane, potentially explaining the discrepancy compared to the other replicates. The non-canonical Phe84 conformation was the most favoured throughout the entirety of the replicate during which the butane remained bound. These data support a model in which Phe84 can adopt a non-canonical rotamer only when stabilised by additional groove-occupying moieties, suggesting a conditional gating function that is responsive to the lipid microenvironment.

## Discussion

Our study reveals a previously unrecognised mechanism of ligand-sensitive structural plasticity in human CD1d, centred on a conserved aromatic residue, Phe84, that acts as a conditional gate over the F′ groove (10). High-resolution crystallography combined with MD simulations and comparative analysis across CD1 and MHC molecules indicates that Phe84 is conformationally flexible, adopting ligand-dependent rotameric states which modulate F′ groove closure and thereby influence both lipid encapsulation and TCR docking. This behaviour is reminiscent of conformationally permissive gating seen at position 84 of MHC class I molecules (24, 27), suggesting that CD1d exploits a similar strategy to accommodate antigen diversity.

Sequence analysis further shows that Phe84 is conserved across most CD1d-expressing species except rodents, which encode a non-aromatic residue at this position (31). These species-specific differences highlight a fundamental divergence between rodent and primate CD1d and caution against over-interpretation of murine models when studying CD1d-restricted T cells. Importantly, while conformational plasticity has been observed in other CD1 isoforms (7, 9), our data provide the first direct crystallographic evidence for rotameric switching in human CD1d itself.

Beyond side-chain motion, groove adaptability may involve subtle backbone shifts, as indicated by our α-helix overlays. Consistent with this, computational analyses have reported extensive helix mobility within the CD1d groove, with conformational dynamics in their study driven by pH-sensitive Trp residues rather than the Phe84 gating observed here. Together, these complementary findings underscore that groove remodelling involves multiple, residue-specific mechanisms of plasticity that enable CD1d to accommodate structurally diverse lipids (32). Conservation of similar aromatic gating residues in classical HLA-I molecules (e.g., Tyr84), non-classical HLA-E, HLA-G and Qa-1, and even in viral MHC-I mimics such as UL18 (22–24, 27, 33) implies that such gating elements constitute a more general evolutionary solution to the challenge of antigen diversity.

Interestingly, CD1d also binds chemically diverse lipids including small sulphur-containing compounds (21, 34) and triacylglycerols (TAGs) (20), unlike the preferential hydrocarbon ligands of CD1b and CD1c. Ligand-stabilised conformational flexibility at Phe84 may therefore provide a structural basis for accommodating bulky species such as PPBF and TAGs by enabling partial groove opening.

In summary, our structures demonstrate that Phe84 operates as a ligand-sensitive aromatic switch in human CD1d, providing a conserved, evolutionarily honed mechanism of structural adaptability that broadens lipid antigen discrimination and validates macaques as relevant models for CD1d-restricted immunity.

## Materials and Methods

### Generation of soluble CD1d proteins

Plasmids encoding the extracellular domains of human CD1d (26), rhesus macaque CD1d, and human β2m were separately cloned into the prokaryotic expression vector pET23d (Novagen), and recombinant proteins were generated separately as inclusion bodies in Escherichia coli Rosetta strain (Novagen). Inclusion bodies were thoroughly washed and fully denatured and reduced in 6 M guanidine-HCl and 20 mM DTT before *in vitro* refolding. Refolding of human and macaque CD1d/β2m was carried out by oxidative in vitro refolding as previously described (26, 35), in the presence of α-GalCer (Avanti Polar Lipids). Prior to refolding, α-GalCer was solubilised in vehicle containing 8.7 mg/ml NaCl and 0.5% Tween-20. Correctly folded proteins were purified by repeated FPLC (Pharmacia) size-exclusion chromatography using preparative grade SD75 26/60 and analytical grade SD75 GL 10/300 gel filtration columns (GE Healthcare).

### Protein Crystallography

Proteins (in buffer 20 mM TrisHCl, pH 7.5, and 50 mM NaCl) were crystallised using sitting-drop vapour diffusion set up in 96 well plates at 20 °C. Human CD1d was set up at 10.5 mg/ml using an Oryx 8 crystallization robot (Douglas Instruments) and crystallised in conditions C12 and G12 (12.5% w/v PEG 1000, 12.5% w/v PEG 3350, 12.5% v/v MPD, 0.1 M bicine/Trizma base, pH 8.5; additives in C12: 0.03 M sodium nitrate, 0.03 M disodium hydrogen phosphate, 0.03 M ammonium sulfate; additives in G12: 0.02 M sodium formate, 0.02 M ammonium acetate, 0.02 M trisodium citrate, 0.02 M sodium potassium l-tartrate, 0.02 M sodium oxamate) from the Morpheus crystallisation screen (Molecular Dimensions) (36). A crystal from the condition G12 was used for the diffraction experiment. Macaque CD1d was set up at 9.5 mg/ml and crystallised in condition A2 of the Morpheus crystallisation screen (10% w/v PEG 8000, 20% v/v ethylene glycol, 0.03 M magnesium chloride, 0.03 M calcium chloride, 0.1 M MES/imidazole, pH 6.5). Crystals were harvested in their mother liquor and flash cooled in liquid nitrogen. Diffraction data collections from Human and Macaque CD1d crystals were performed at the European Synchrotron Radiation Facility (ESRF), beamline ID30B, France (37), and the Diamond Light Source, beamline I03, UK, respectively. Data were processed in *xia2* (38) using *DIALS* [(39, 40) at 1.76 Å and 1.83 Å resolution (Table 1). Molecular replacement was carried out with *MOLREP* (41) from the *CCP4 suite* (42), using *AlphaFold2* (43, 44)predictions as search models. Iterative model building with *Coot* (45) and refinement against structure factor amplitudes with *Refmacat*/*Refmac5* (46) and against intensities with *Servalcat* (47) yielded the final structure models *R*_work_/*R*_free_ values of 0.2132/0.2542 and 0.1707/0.2169, respectively (Table 1). The structures were validated and deposited in the Protein Data Bank (33) under the accession codes 9RSE and 9RSF. The diffraction images are available at Zenodo at https://doi.org/10.5281/zenodo.15442781 and https://doi.org/10.5281/zenodo.15441697.

### iNKT cells

Blood samples were obtained from healthy local volunteers after informed consent. Peripheral blood mononuclear cells (PBMCs) were isolated from human peripheral venous blood by density gradient centrifugation (Ficoll-Paque PLUS, GE Healthcare, Amersham, UK). iNKT cell lines were generated from PBMCs by seeding at 2 × 10^6^ cells/ml into 24 well plates in 1 ml iNKT cell growth medium containing RPMI 1640 (Lonza, Slough, UK), 10% foetal bovine serum (FBS; Sigma), penicillin (100 IU/ml)/streptomycin (100 μL/ml; Gibco, Thermo Fisher Scientific, Basingstoke, UK), 1% non-essential and essential amino acids, 1% L-glutamax (both Sigma), 55 µM 2-mercaptoethanol (Fisher Scientific, Loughborough, UK) and 15 mM HEPES (Gibco). Before use, the solution was filtered (Stericup Quick Release Durapore 0.22 μm PVDF, Millipore, Merck group, London, UK). PBMCs were pulsed with α-GalCer (100 ng/ml) on day 0 and rhIL-2 (200 IU/ml; Proleukin, Chiron) added on day 7. Cells were fed every 2-3 days with fresh iNKT cell growth medium containing rhIL-2 (200 IU/ml). Following the 14-day expansion live CD3+, CD1d-α-GalCer tetramer+, Vβ11+ T cells were bulk sorted into tubes containing iNKT cell growth medium. Sorted lines were stimulated with 1 µg/ml phytohemagglutinin (PHA-L; Remel), rhIL-2 (200 IU/ml) in the presence of 2 × 10^6^ /ml autologous γ-irradiated (50 Gy) PBMCs. Following a two-week expansion pure iNKT cell lines were utilised for tetramer staining and for functional assays.

### Flow cytometry

The following fluorescent reagents were used: PE-conjugated human or macaque CD1d tetramers loaded with α-GalCer, FITC-conjugated anti-human TCR Vα24 (clone C15; Beckman Coulter Ltd, High Wycombe, UK), FITC-conjugated anti-human TCR Vβ11 (clone C21; Beckman Coulter Ltd), BV421-conjugated anti-human CD3 antibody (clone UCHT1; Biolegend), PE-conjugated anti-CD69 (clone FN50), PE/Cy7-conjugated anti-CD25 (clone BC96) (Both Biolegend), and propidium iodide for live/dead staining (Sigma). Cells were stained with mAbs in phosphate-buffered saline containing 1% FBS and 1 mM EDTA for 30 minutes at 4°C, acquired on BD FACS Aria (BD Biosciences) and analysed on FlowJo V10.9.0 software (FlowJo LLC, Oregon, USA).

### CD1 tetramers

Refolded CD1d complexes were biotinylated via an engineered BirA motif at the C terminus, repurified by size exclusion chromatography, and used to generate fluorescently labelled CD1 tetramers by conjugating them to phycoerythrin (PE)–streptavidin (Sigma).

### Cytokine ELISA

iNKT cells, 1 × 10^5^ per well, were stimulated for 24 hours in 96-well plates that were pre-bound with different concentrations of human or macaque CD1d-α-GalCer complexes. Culture supernatants were then analysed for cytokine concentrations for IL-2 using an in-house developed ELISA kit.

### Surface Plasmon Resonance

Streptavidin (∼5,000 RU) was amino-coupled to a Biacore CM-5 chip (BIAcore AB) and 50 μg/mL biotinylated lipid-CD1d complexes or control protein CD1c-endo were loaded on individual flow cells until the response measured ∼1,000 RU. Recombinant iNKT cell TCRs were serially diluted and flowed over the protein-loaded flow cells at a rate of 5 or 50 μL/min for determination of equilibrium binding or kinetics. Responses were recorded in real time on a Biacore 3000 machine at 25 °C, and data were analysed using the BIA evaluation software (GE Healthcare, Buckinghamshire, U.K.) as described previously (25).

### Soluble iNKT cell TCRs

iNKT TCR heterodimers were generated as previously described (25, 48). Briefly, the extracellular region of each TCR was produced separately from *Escherichia coli* Rosetta (DE3) pLysS competent cells (Novagen). To produce stable, disulfide-linked heterodimers, cysteines were incorporated into the TCR α- and β-chain constant regions, replacing residues Thr48 and Ser57, respectively. Expression, refolding, and purification of the disulfide-linked iNKT cell TCR αβ heterodimers were carried out as previously described (25). Purified refolded TCR proteins were assessed by both reducing and nonreducing SDS/PAGE analysis.

### Molecular dynamics simulations

Molecular dynamics simulations were performed using Gromacs1 2021.4 patched with Plumed2 v2.7.3. Systems were protonated using the H++ webserver (49) to a target pH of 7.0. For REST2 MD simulations proteins were modelled using the Amber19SB forcefield and α-GalCer was modelled using the GAFF2 (50) forcefield. Systems were solvated up to 10 Å beyond the protein with TIP3P (51, 52) waters and neutralised using additional sodium ions. Each system was minimised for 5000 steps using steepest descent minimization. Periodic boundary conditions were applied, and hydrogen bonds were constrained using the LINCS algorithm. Non-bonded interactions were cut-off at 1.2 nm, with an additional force-switching function implemented at 1.1 Å for van der Waals interactions. After minimisation, systems were equilibrated to 300 K in the NVT ensemble for 100 ps using the velocity-rescaling thermostat (53) with 0.1 ps time constant and 2 fs timestep. Additional long range dispersion corrections were applied. Additional equilibration was performed in the NPT ensemble at 1 bar pressure with the Berendsen barostat and 2.0 ps time constant. The final production simulation was performed in the canonical NVT ensemble with the nose-hoover thermostat (2.0 ps time constant). Replica Exchange with Solute Scaling (REST2) (54) was performed using 18 replicas distributed between effective temperatures of 300-450 K. Simulations were performed for approximately 200 ns per replica. Scaling was applied to all atoms in the lipid binding domain (α1 and α2 domains) and α-GalCer. Simulation frames were recorded every 10 ps. Conventional MD simulations with butane were performed for 500 ns per replica.

### Statistical Analysis

GraphPad Prism version 8.00 (GraphPad Software, Inc.) was used for statistical analysis, and P values ≤0.05 were considered statistically significant. The Mann–Whitney U test, and T test, were used as stated in the figure legends.

## Acknowledgements

We thank Carolann McGuire and Sarah Pearson for their assistance with flow cytometry (FACS facility, Faculty of Medicine, University of Southampton). We thank Chris Holes for support with macromolecular crystallisation. We are especially indebted to Regina Teo for her expert laboratory management and sustained support throughout the project. D.B was supported by a studentship funded by the Institute for Life Sciences and the Faculty of Medicine, University of Southampton. A.L was supported by a MRC studentship (MR/W007045/1). M.M. was supported by the BBSRC (grant No.BB/Y008839/1), the STFC (grant No. 8521412). P.E was supported by MRC (MR/P023754/1 and MR/W025728/1). S.M was supported by MRC (MR/S024220/1) and Cancer Research UK (23562). We thank the Southampton National Institute for Health Research Biomedical Research Centre for infrastructure support.

---

**Supp Fig. 1.**
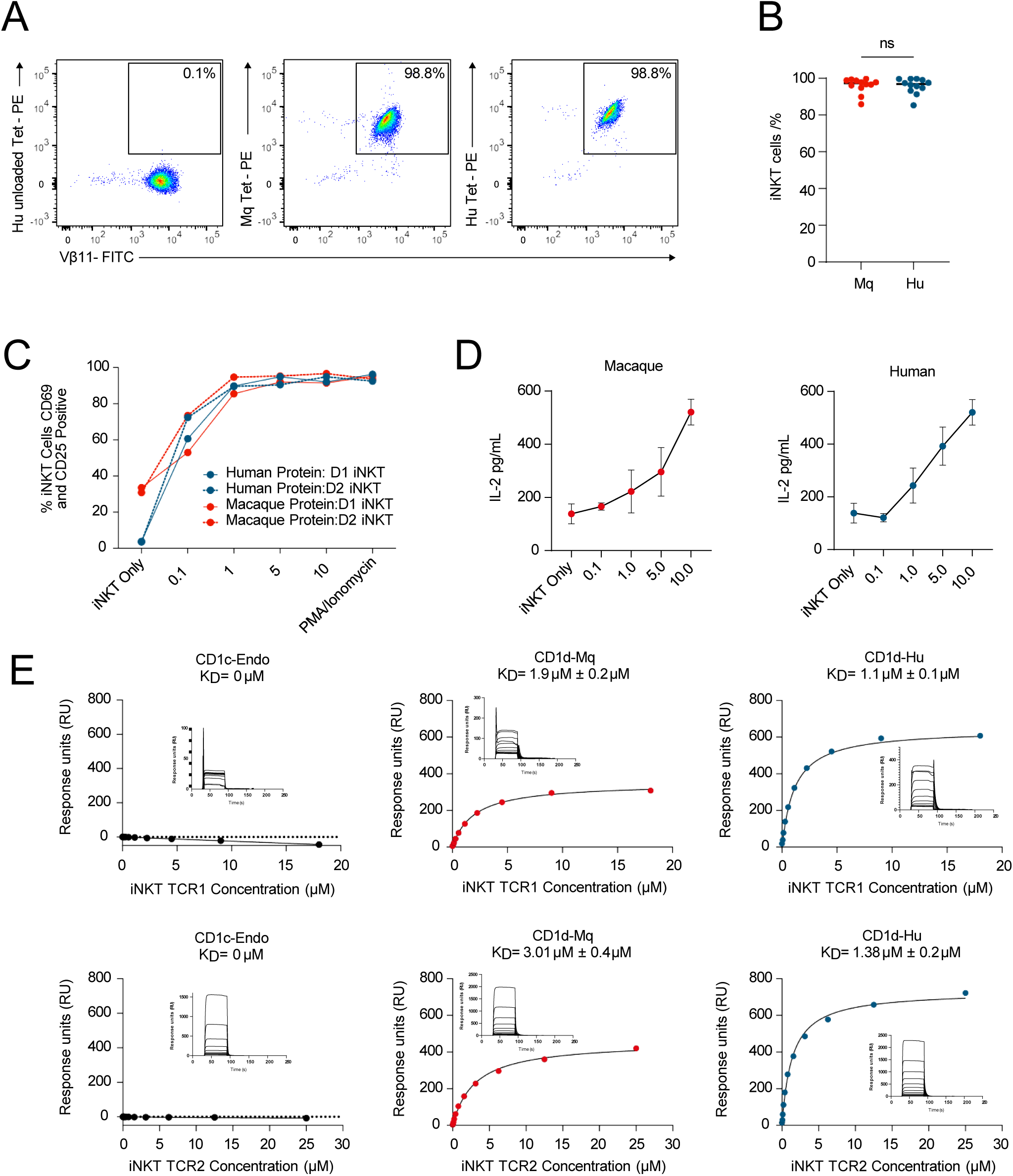
Macaque and human CD1d activate iNKT cells and bind iNKT cell TCRs. **A)** Representative flow cytometry plots showing the staining of human iNKT cells by macaque and human CD1d-tetramers. Live CD3^+^CD1d-tetramer^+^Vβ11^+^ cells are shown. Plots show parentage of iNKT cells. Pure human iNKT cell lines were generated by *in vitro* expansion and flow cytometry guided cell sorting. Human CD1d-endo-tetramer was used as control. **B)** Cumulative data showing the percentage of iNKTs stained by human and macaque CD1d tetramers. iNKT cell lines from 11 healthy donors were stained. **C)** Plate bound activity assay showing the specific activation of iNKT cells in response to macaque and human CD1d in a dose dependent manner. iNKT lines from two donors, D1 and D2, were used. **D)** IL-2 secretion by human iNKT cells cultured with macaque and human CD1d. **E)** Surface plasmon resonance measurements (BiaCore) for binding of 2 human iNKT-cell TCRs to immobilised CD1c-endo (Left), macaque CD1d-α-GalCer (middle), and human CD1d-α-GalCer (right) complexes at equilibrium. Kd, calculated dissociation constant; RU, response units. Data in E are representative of two independent experiments.

**Supplementary Fig. 2.**
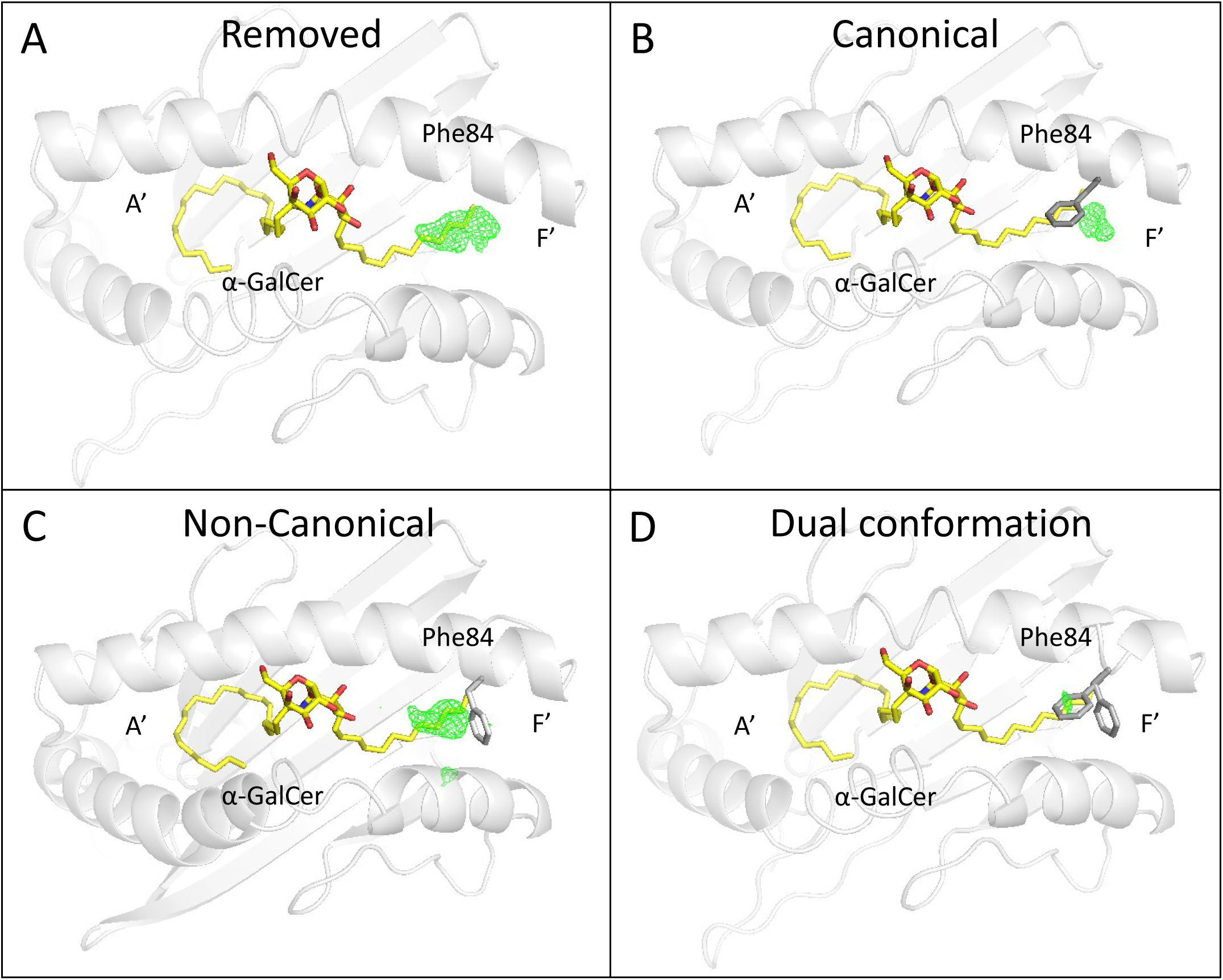
Electron density supports dual Phe84 conformations. Flexible Phe84 residue shown in different orientations using omit maps with resulting difference density following refinement, contoured at 3σ and coloured in green. The residue is shown **A)** omitted, **B)** canonically placed, **C)** non-canonically placed, and **D)** in dual conformation. Human CD1d, grey; α-GalCer, yellow (coloured by heteroatom with nitrogen in blue and oxygen in red). Phe84 is shown in stick representation. All individual images were generated using Pymol.

**Supplementary Fig. 3.**
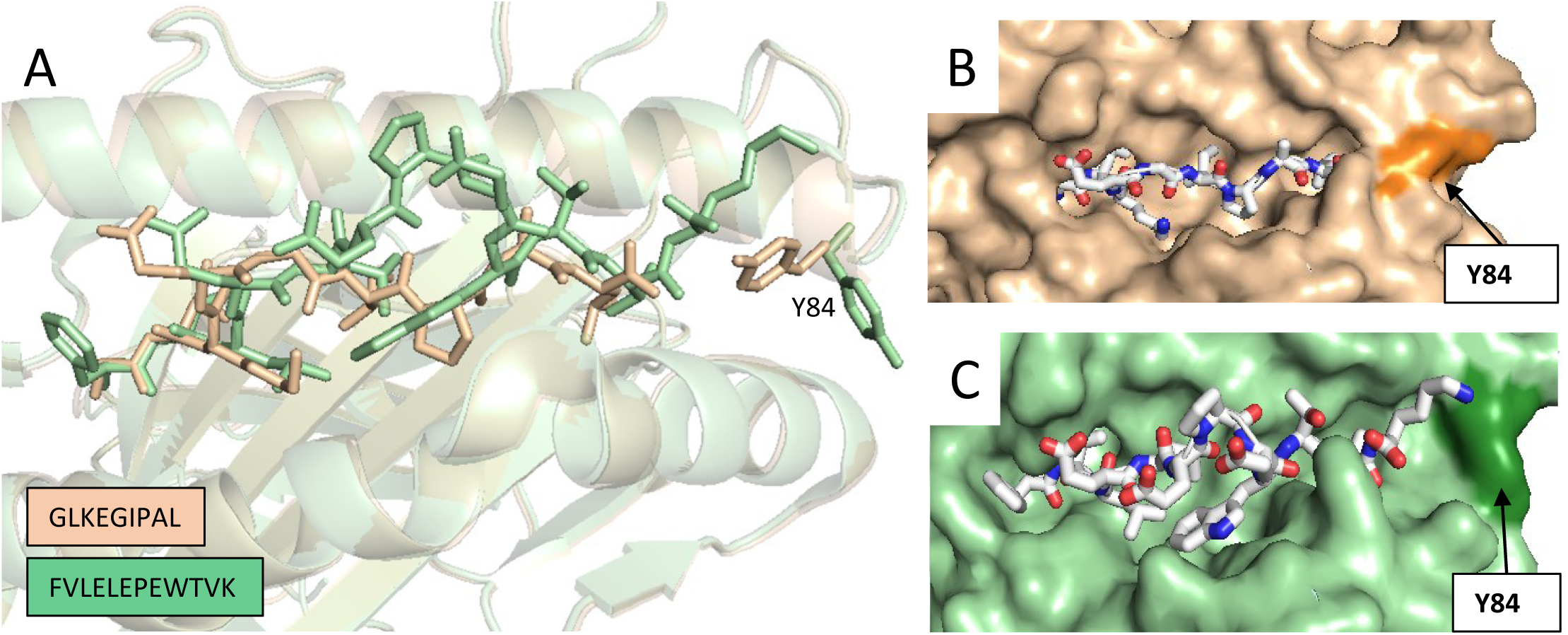
Conformational flexibility at the HLA-A*02:01 Fʹ groove remodels the peptide binding groove. **A)** Superposition of HLA-A02:01 bound to a canonical “nested” 9-mer peptide (GLKEGIPAL, gold; PDB: 5ENW) and an 11-mer peptide (FVLELEPEWTVK, green; PDB: 5DDH) reveals marked differences at the C-terminal end of the peptide binding cleft. **B)** Surface representation of HLA-A02:01-GLKEGIPAL, highlighting the closed conformation of Tyr84, which caps the Fʹ-pocket and accommodates shorter peptides in a buried, canonical manner. **C)** In contrast, binding of the extended 11-mer peptide to HLA-A*02:01 results in rotation of Tyr84 by ∼90°, opening the Fʹ-pocket and allowing protrusion of the peptide C-terminus from the groove. This illustrates the structural plasticity of MHC class I molecules in accommodating non-canonical, extended peptide ligands.

**Supplementary Fig. 4.**
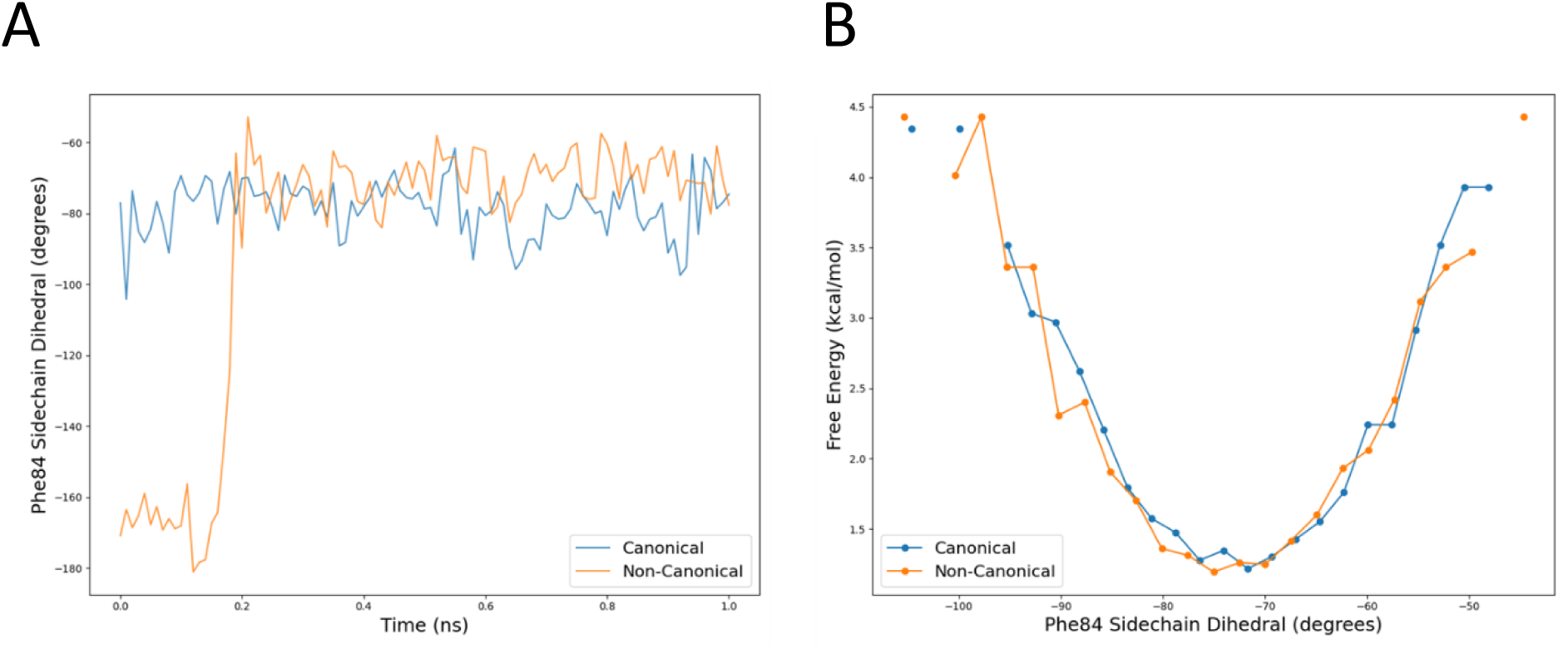
MD simulations confirm canonical Phe84 is energetically favoured in absence of ligand. **A)** Dihedral angle of Phe84 during NVT equilibration from both canonical and non-canonical starting states. **B)** Potential of mean force for Phe84 sidechain rotation from REST2 simulations, showing energetic favourability of canonical rotamer unless stabilised by groove-occupying ligand.

